# Development of a novel PROTAC using the nucleic acid aptamer as a targeting ligand for tumor selective degradation of nucleolin

**DOI:** 10.1101/2021.08.05.455211

**Authors:** Lin Zhang, Ling Li, Xia Wang, Huimin Liu, Yibin Zhang, Tiantian Xie, Hui Zhang, Xiaodong Li, Tianhuan Peng, Xing Sun, Jing Dai, Jing Liu, Wencan Wu, Mao Ye, Weihong Tan

**Author notes:** Correspondence: Mao Ye, Molecular Science and Biomedicine Laboratory (MBL), State Key Laboratory of Chemo/Biosensing and Chemometrics, College of Biology, College of Chemistry and Chemical Engineering, Aptamer Engineering Center of Hunan Province, Hunan University, Changsha, Hunan 410082, China, Correspondence: Weihong Tan, The Cancer Hospital of the University of Chinese Academy of Sciences (Zhejiang Cancer Hospital), Institute of Basic Medicine and Cancer (IBMC), Chinese Academy of Sciences, Hangzhou, Zhejiang 310022, China. These authors contributed equally to this work (Lin Zhang, Ling Li, Xia Wang).

## Abstract

PROteolysis TArgeting Chimeras (PROTACs) induce targeted protein degradation by hijacking the intracellular ubiquitin proteasome system, thus emerging as a new strategy for drug development. However, most PROTACs generated lack cell-type selectivity and have poorly soluble in water. To address this drawback, we developed a novel PROTAC ZL216 using aptamer AS1411 as a targeting ligand of nucleolin to conjugate with a small molecule ligand of E3 ligase VHL, which shows high aqueous solubility and serum stability. Based on the differential expression of nucleolin on the cell surface, ZL216 could bind to and internalize into breast cancer cells, but not normal breast cells. Furthermore, we revealed that ZL216 promoted the formation of a nucleolin-ZL216-VHL ternary complex in breast cancer cells and potently induced nucleolin degradation *in vitro* and *in vivo*. As a result, ZL216 inhibited the proliferation and migration of breast cancer cells. These studies demonstrate that in addition to peptides and small molecule compounds, nuclei acid aptamers can also be used to generate PROTACs, which broadens the toolbox constructing PROTACs and provides a promising strategy for development of tumor-selective PROTACs.

## Introduction

PROteolysis Targeting Chimeras (PROTACs) are small molecules with heterobifuntional property, consisting of a ligand for binding to a protein of interest (POI) and the other for recruiting an E3 ubiquitin ligase via a linker connection.(1, 2) Such molecules can tether an E3 ligase to POI by forming a ternary complex, thereby triggering proximity-induced ubiquitination and degradation of POI in a catalytic manner.(3) PROTACs have been developed to degrade proteins that play important roles in tumorigenesis and do so with extraordinary efficacy on multiple tumor types, including BRAF mutant V600E for melanoma,(4) BET for prostate cancer,(5) BCL-XL for leukemia(6) and Estrogen Receptor (ER) for breast cancer.(7) Until present, at least 15 targeted protein degraders are expected to be tested in patients at different stages of human clinical trials.(8) However, although PROTACs are highly efficient for protein degradation and, hence, a potential therapeutic strategy for cancer, most PROTACs generated lack tumor selectivity, one of the main reasons is that the E3 ligases utilized by PROTACs are widely expressed in both normal and tumor tissues. As a result, these PROTACs may produce on-target toxicity if POIs are not tumor-specific. This calls for the development of selective ligands able to restrict the activity of PROTACs to tumor cells, but not normal cells, thus generating tumor-selective PROTACs. To overcome limitation of PROTACs selectivity, antibody-based PROTACs were recently developed, which utilized antibody as targeting elements to achieve cell-specific delivery of PROTAC and degradation of POIs. (9–12)

Aptamers are single-stranded DNA or RNA oligonucleotides with a length typically less than 100 nucleotides.(13) Aptamers are generated by an iterative selection process termed systematic evolution of ligands by exponential enrichment (SELEX) from a randomized oligonucleotide library.(14) Similar to antibodies, aptamers bind to their targets with high specificity and affinity by folding into a unique three-dimensional conformation. Compared to protein antibodies, aptamers hold unique characteristics derived from their oligonucleotide properties, such as low immunogenicity and toxicity, easy chemical synthesis and modification, rapid tissue penetration and excellent stability.(15) With the advent of cell-specific aptamers, aptamers have been widely used as tumor recognition ligands for targeted therapy and diagnostics. For example, CD71 ssDNA aptamer was coupled with doxorubicin to specifically target tumor cells and inhibit their proliferation.(16)

AS1411 is a 26-base guanine-rich ssDNA aptamer, which binds to nucleolin with high affinity and specificity. Many studies have indicated that enhanced expression of nucleolin on the cell surface is restricted to tumor cells, but not normal cells, thus conferring a tumor-selective binding behavior to AS1411.(17) Meanwhile, AS1411 can be internalized into tumor cells by nucleolin-dependent macropinocytosis and subsequently exerts antiproliferative activity against a wide range of tumor types, including breast cancer,(18) gloma,(19) renal cell carcinoma(20) and leukemia.(21) Thus, AS1411 potentially acts as both a tumor-targeting element and an anticancer therapeutic agent. Recently, AS1411 was used as a targeted delivery tool to deliver BET-PROTAC into breast cancer cells for degrading BRD4 protein.(22)

In this study, we report the first proof-of-concept evidence using nucleic acid aptamer to construct a novel PROTAC. Aptamer AS1411, as a targeting ligand for nucleolin, was conjugated to a small molecule ligand of E3 ligase VHL via a DBCO-azide click reaction, which generated a nucleolin-targeting PROTAC ZL216 with excellent serum stability and water solubility. We demonstrated that ZL216 could recruit E3 ligase VHL to nucleolin in breast cancer cells and potently induce the degradation of nucleolin *in vitro* and *in vivo*. As a result, ZL216 inhibited the proliferation and migration of breast cancer cells. Our work demonstrates the potential use of aptamers in generating PROTACs to achieve selective degradation of target protein in tumor cells and provides a promising strategy for development of tumor-selective PROTACs.

## RESULTS

### Development of a novel PROTAC ZL216 using aptamer

By analyzing various cancer types found on the TCGA database using TIMER2.0, we found that the gene expression levels of nucleolin in multiple tumor tissues, including breast cancer tissues, were significantly higher than those of normal tissues (Figure S1A). Consistent with this, the result of the CPTAC proteomics database also showed elevated protein levels of nucleolin in breast cancer tissues compared to corresponding normal tissues (Figure S1B). We further confirmed this finding in cell lines by Western blotting, which showed enhanced expression of nucleolin protein in breast cancer cells (MCF-7 and BT474), but not in immortalized breast epithelial cells (MCF-10A) (Figure S1C). Moreover, a part of nucleolin was distributed at the surface of MCF-7 and BT474 cells, but not the surface of MCF-10A cells (Figure S1D), which is consistent with the notion that the expression of nucleolin on the cell surface is restricted to tumor cells.

AS1411 functions as an ssDNA aptamer that can bind to nucleolin with high affinity and specificity and that it is quickly internalized into cells following binding to nucleolin.(17) Given the critical role of nucleolin in supporting tumorigenesis and metastasis, we hypothesized that AS1411-constructed PROTAC could achieve targeted antitumor activity through promoting nucleolin degradation specifically in tumor cells (Figure 1A). To test the possibility, DBCO-labeled AS1411, as a targeting element for nucleolin, was conjugated via the DBCO-azide cycloaddition reaction to a VHL E3 ligase-binding ligand, (S, R, S)-AHPC-PEG3-azide (AHPC), to generate a nucleolin degrader ZL216 (Figure 1B and 1C), whereas an ssDNA sequence, which has no affinity for nucleolin, was linked to AHPC as a negative control (Figure S2). Similarly, the synthesis and characterization of Cy5-labeled Control and ZL216 were shown in Figure S3. The purified products were also identified by DNA mass spectrometry (Figure S4).

**Figure 1.**
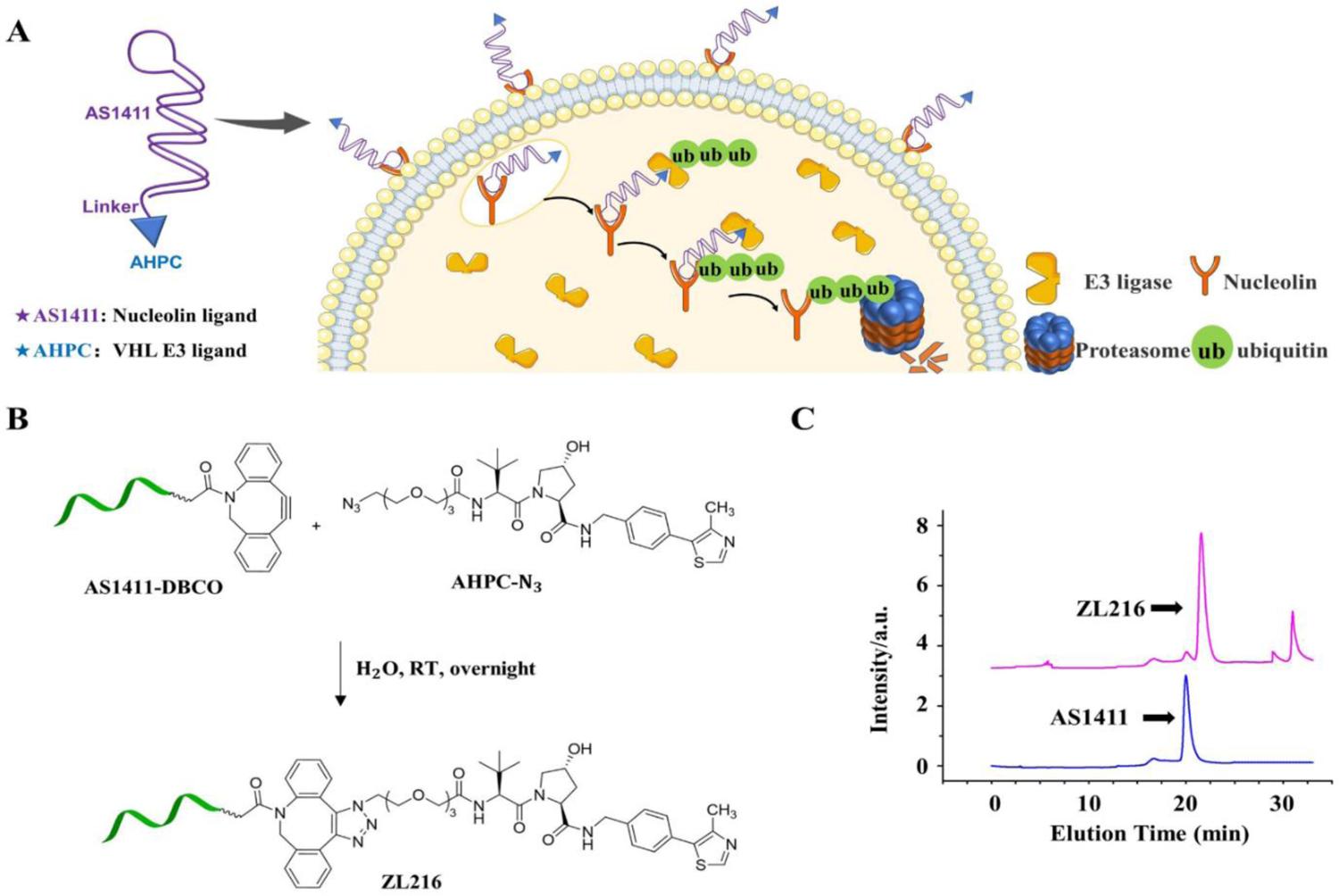
Development of aptamer PROTAC ZL216. (A) Presumed mode of action of ZL216 on degradation of nucleolin in cancer cell. (B) Synthetic roadmap and LC-MS analysis of ZL216. (C) The purification of ZL216 by HPLC.

To measure the stability of ZL216 in serum, ZL216 and AS1411 were incubated with human serum. We found that ZL216 held an excellent serum stability, similar to that of AS1411, with a half-life is 70.5 hours (Figure S5). Meanwhile, the water solubility of ZL216 was significantly enhanced compared to that of free VHL E3 ligase-binding ligand AHPC through conjugation with aptamer AS1411 (Figure S6).

### Selective binding ability of ZL216

Previous studies have shown that AS1411 can specifically bind to breast cancer cells through recognizing cell-surface nucleolin.(23, 24) To assess whether ZL216 still retains the binding specificity to breast cancer cells, cells were incubated with Cy5-labeled AS1411, ZL216 and Controls, respectively. As shown in Figure 2A, 2B and 2C, a significant fluorescence peak shift was observed from MCF-7 and BT474, but not MCF-10A cells, indicating that ZL216 held its binding specificity to breast cancer cells in a manner similar to that of AS1411. Meanwhile, the selective binding ability of ZL216 was further confirmed by confocal imaging (Figure 2D, 2E and 2F). To identify whether ZL216 has the same binding sites as AS1411, a competitive binding assay was performed by flow cytometry. Different concentrations of unlabeled AS1411 were incubated with Cy5-labeled ZL216. Intriguingly, ZL216 gradually lost its binding to MCF-7 and BT474 cells as the concentration of AS1411 increased, whereas the Control showed no effect from the binding between ZL216 and MCF-7 or BT474 cells, implying that the same binding sites were shared by ZL216 and AS1411(Figure 2G and 2H). Furthermore, the binding affinity assay showed that the K_d_ of ZL216 for MCF-7 and BT474 cells was 11.92 ± 3.7 nM and 15.27 ± 2.8 nM, while the K_d_ of AS1411 on MCF-7 and BT474 cells was 13.93 ± 3.6 nM and 30.76 ± 7 nM, respectively (Figure S7), suggesting that ZL216 held the excellent binding ability to breast cancer cells.

**Figure 2.**
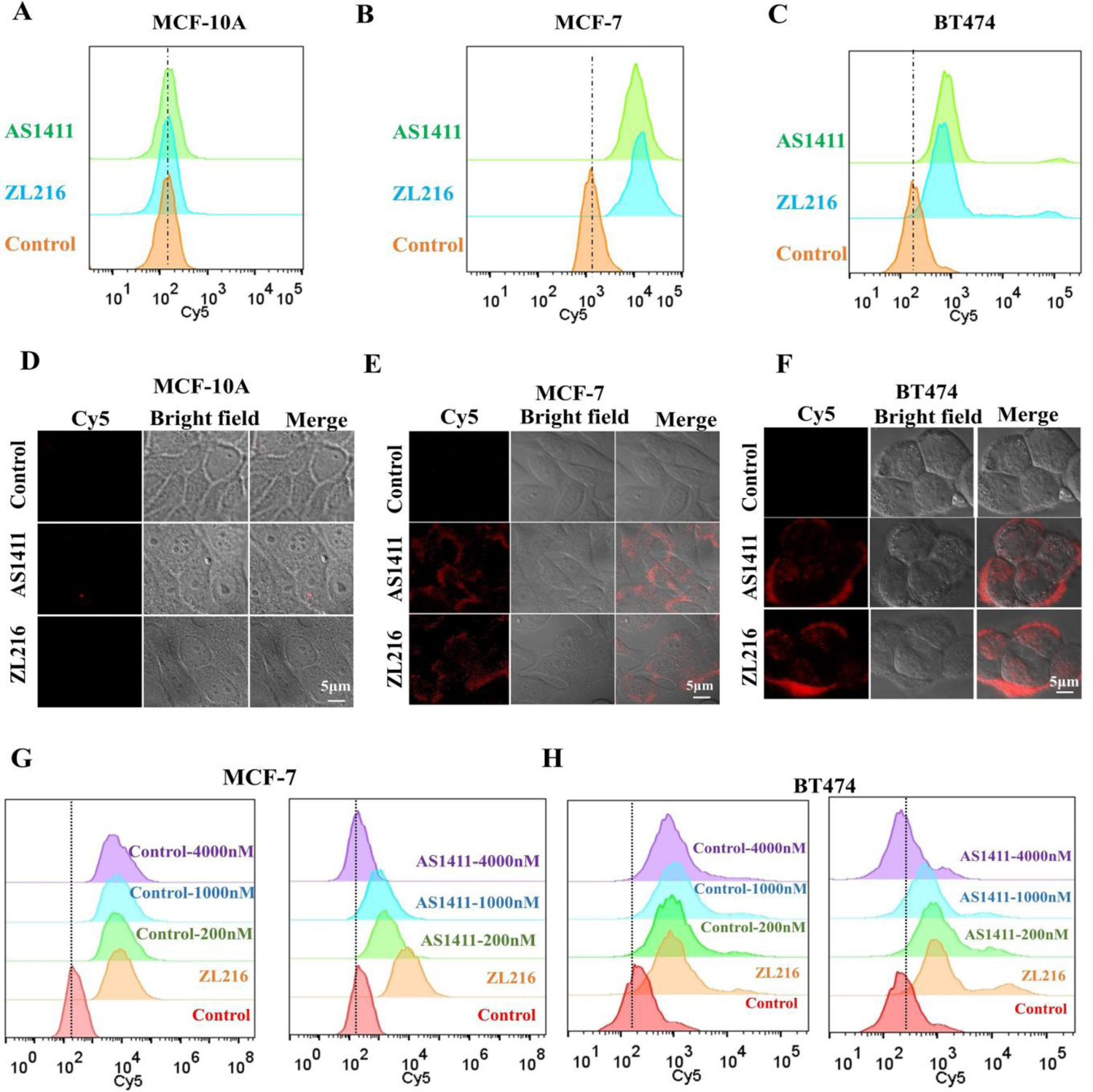
ZL216 selectively binds to breast cancer cells. (A, B, C) The binding of ZL216 and AS1411 to MCF-10A (A), MCF-7 (B) and BT474 (C) cells was analyzed by flow cytometry. The concentrations of ZL216 were 25 nM in MCF-7 or 50 nM in BT474 and MCF-10A cells. (D, E, F) MCF-10A (D), MCF-7 (E) and BT474 (F) cells were incubated with Cy5-labeled ZL216 and AS1411. Fluorescence, bright-field, and merged confocal images were shown. (G, H) MCF-7 (G) and BT474 (H) cells were incubated with the indicated concentration of AS1411, followed by treatment with Cy5-ZL216. Competition binding was analyzed by flow cytometry.

### ZL216 specifically attenuates nucleolin of breast cancer cells *in vitro* and *in vivo*

Internalization into cells directly determines whether aptamer-constructed PROTAC can hijack the intracellular ubiquitin proteasome system. To investigate the internalization property of ZL216, intracellular fluorescence signals were detected by flow cytometry after eliminating the fluorescence signals on the cell surface by trypsin or proteinase K. As shown in Figure S8, the removal of the fluorescence signals on the cell surface reduced, rather than eliminated signals from target cells, indicating that a fraction of ZL216 could be internalized into target cells. Next, we evaluated ZL216 for its ability to regulate nucleolin levels. After cells were treated with different concentrations of ZL216, nucleolin abundance was analyzed by Western blotting. We found that ZL216 could effectively downregulate nucleolin levels in MCF-7 and BT474 cells, but not MCF-10A cells (Figure 3A, 3B and 3C). At the 5-h time point, the half-maximal inhibition concentration (DC_50_) and maximum inhibition (D_max_) of MCF-7 cells were 13.5 nM and 77%, while those of BT474 cells were 18.4 nM and 60%, respectively. The time dependency of ZL216 on nucleolin showed that maximal reduction of nucleolin was achieved at 48 h for MCF-7 cells and 5 h for BT474 cells. Meanwhile, the degradation of nucleolin was not detected in MCF-10A cells (Figure 3D, 3E and 3F). In addition, both AS1411 and an E3 ligase-binding ligand AHPC had no effect on nucleolin levels (Figure S9). Furthermore, to determine whether ZL216 retains it recognition ability and reduce nucleolin levels *in vivo*, we generated MCF-7 and BT474 xenografts in mice and then treated the mice with Cy5-labeled ZL216, AS1411 and Control via tail vein injection. As expected, the fluorescence signal was rapidly accumulated at the tumor sites after injecting ZL216 and AS1411, while no significant fluorescence signal was observed at the tumor sites treated with Control, indicating that ZL216 held the same ability to target tumor *in vivo* as that achieved by AS1411 (Figure 3G and 3H). Notably, ZL216 could decrease nucleolin levels in xenografts (Figure 3I and 3J). In contrast, neither AS1411 nor AHPC had any effect on nucleolin levels (Figure 3K and 3L). This result highlighted that ZL216 not only possessed an ability to target tumors similar to that of AS1411, but also had the added benefit of reducing nucleolin levels, not otherwise achievable by AS1411.

**Figure 3.**
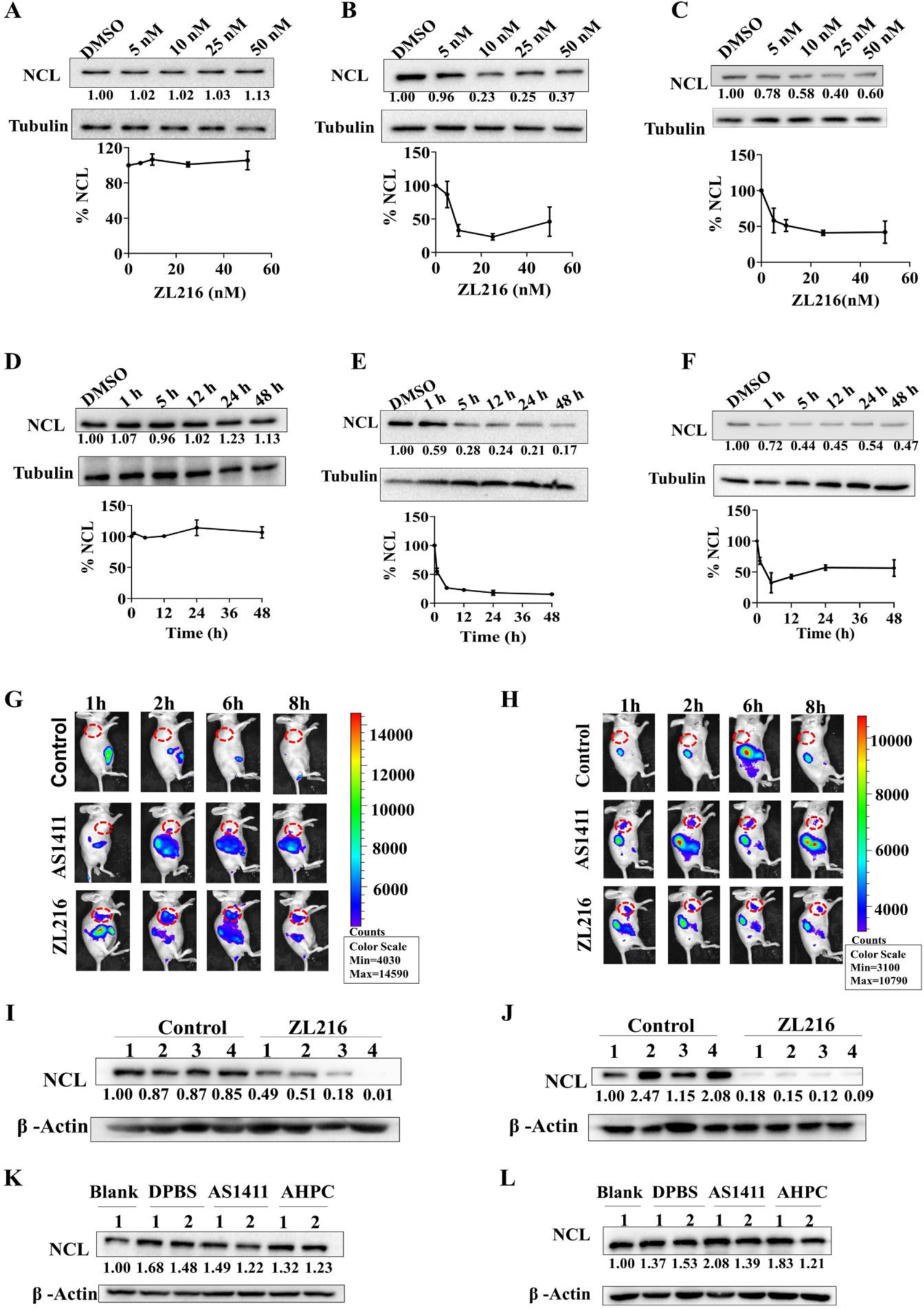
ZL216 downregulates the nucleolin (NCL) of breast cancer cells *in vitro* and *in vivo*. (A,B,C) MCF-10A (A), MCF-7 (B) and BT474 (C) cells were treated with the indicated concentration of ZL216 for 5 h. Total protein was extracted and subjected to Western blotting using the indicated antibodies. (D, E, F) ZL216 treated cells at the concentration of 50 nM for MCF-10A (D), 25 nM for MCF-7 (E) and 50 nM for BT474 (F). Total protein was extracted at the indicated time points and subjected to Western blotting using the indicated antibodies. (G, H) MCF-7 cells (G) and BT474 cells (H) were injected subcutaneously into nude mice. When tumors grew to desired size, 50 μM Cy5-labeled AS1411, ZL216 or Control was injected intravenously into nude mice. The fluorescence images *in vivo* were shown at the indicated time points. (I, J) MCF-7 (I) and BT474 (J) tumor-bearing mice were injected intravenously with 50 μM ZL216 or Control for 8 h. Total protein was extracted from tumor tissues and subjected to Western blotting using the indicated antibodies. (K, L) MCF-7 (K) and BT474 (L) tumor-bearing mice were injected intravenously with 50 μM AS1411, AHPC or DPBS for 8 h. Total protein was extracted from tumor tissues and subjected to Western blotting using the indicated antibodies.

### ZL216 degrades nucleolin in a VHL- and proteasome-dependent manner

To confirm that ZL216 causes the reduction of nucleolin levels through the expected degradation mechanism via proteasome, we first tested the effect of ZL216 on the expression of nucleolin mRNA by real-time PCR. We found that the levels of nucleolin mRNA were not regulated by ZL216, Control, AHPC, AS1411 or DMSO (Figure 4A and 4B), indicating that ZL216 did not affect nucleolin expression at the transcriptional level. Furthermore, downregulation of nucleolin caused by ZL216 could be abolished by a proteasome inhibitor, MG132, or a cullin neddylation inhibitor, MLN4924 (Figure 4C and 4D), suggesting that ZL216 decreased nucleolin levels through the proteasome degradation pathway. To further identify whether ZL216 degrades nucleolin via VHL, cells were treated with a high concentration of free AHPC to compete for the binding of ZL216 to VHL. The results showed that an excess amount of AHPC could significantly attenuate the effect of ZL216 on nucleolin in MCF-7 and BT474 cells (Figure 4E and 4F), demonstrating that ZL216-mediated regulation of nucleolin protein levels depended on VHL. Since ZL216 downregulated nucleolin protein levels, we questioned whether ZL216 affected the stability of nucleolin. To this end, cells were treated with cycloheximide (CHX) to inhibit protein biosynthesis, followed by analyzing nucleolin protein at indicated time points. We found that ZL216 treatment significantly shortened the half-life of nucleolin protein compared with Control treatment (Figure 4G and H).

**Figure 4.**
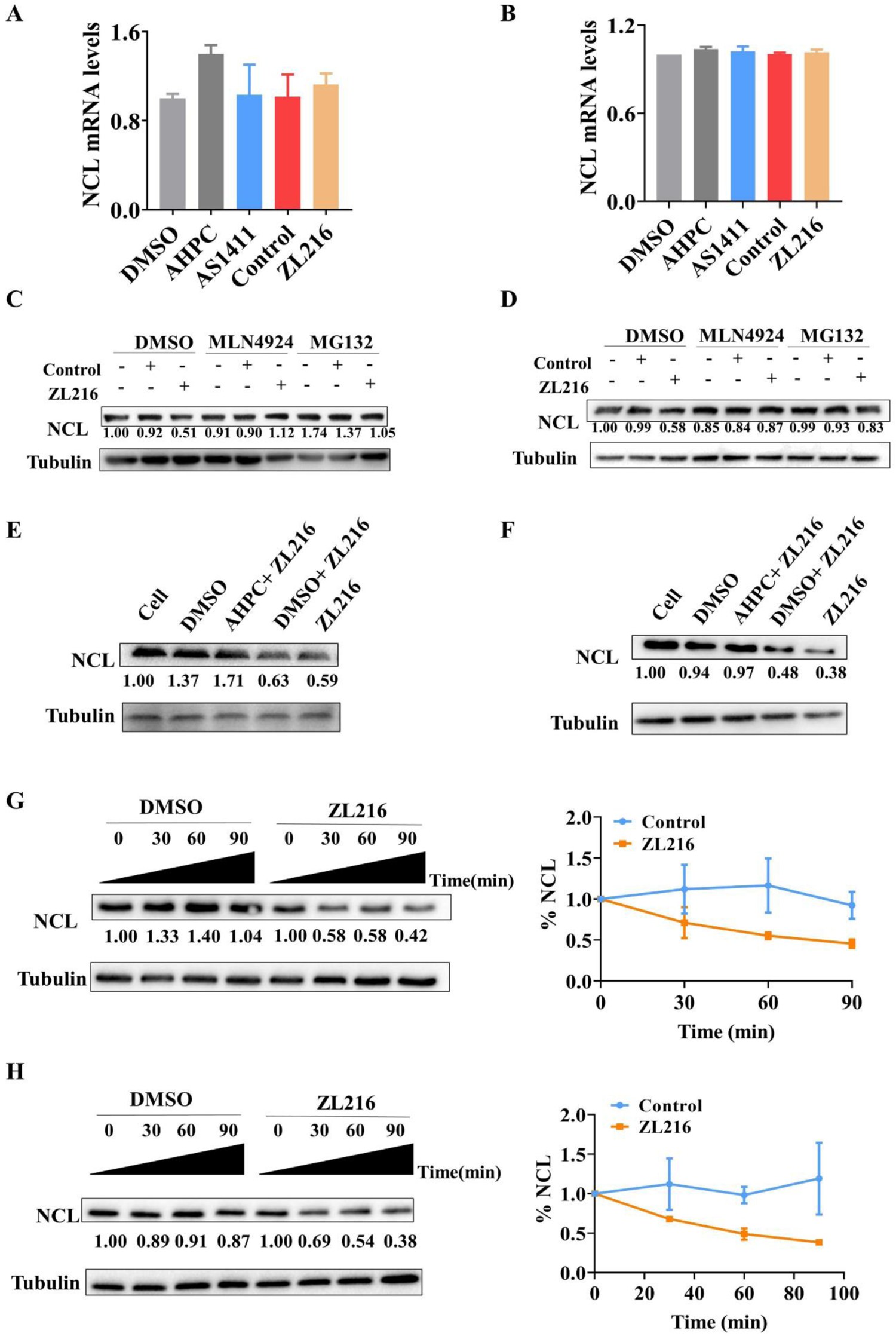
ZL216 redirects E3 ubiquitin ligase VHL to degrade NCL proteins via the proteasomal pathway. (A, B) Total RNA from MCF-7 (A) or BT474 (B) cells treated with Control, AHPC, AS1411 or ZL216 was isolated and subjected to RT-PCR. The error bars represent the SD of triplicate measurements. (C, D) MCF-7 (C) or BT474 (D) cells were pretreated with DMSO, MLN4924 (5 μM) or MG132 (20 μM), followed by ZL216 treatment for 5 h. The indicated proteins were analyzed by Western blotting. (E, F) MCF-7 (E) or BT474 (F) cells were incubated with AHPC (1 μM), followed by ZL216 treatment for 5 h. The indicated proteins were analyzed by Western blotting. (G, H) MCF-7 (G) or BT474 (H) cells were incubated with 100 μg / ml CHX for 1 h, followed by ZL216 treatment. Total proteins were extracted at the indicated time points and subjected to Western blotting using the indicated antibodies. The concentrations of Control, AS1411 or ZL216 were 25 nM for MCF-7 cells or 50 nM for BT474 cells.

### ZL216 induces the formation of nucleolin-ZL216-VHL ternary complex

Nucleolin is not the intracellular natural substrate of the E3 ligase VHL. We hypothesized that ZL216 may mediate the degradation of nucleolin through the formation of a nucleolin-ZL216-VHL ternary complex. To address this, Flag-VHL and Myc-NCL plasmid were transfected into MCF-7 cells, and coimmunoprecipitation was conducted using anti-Flag or anti-Myc antibody. The results indicated that Myc-nucleolin was present in Flag-VHL immunoprecipitates after cells were treated with 25 nM ZL216 and vice versa (Figure 5A and 5B). Conversely, treatment with Control, which lacks nucleolin binding, failed to detect the expected protein in Flag-VHL or Myc-nucleolin immunoprecipitates. Immunofluorescence staining further revealed that ZL216, but not Control treatment, caused the colocalization of nucleolin and VHL (Figure 5C). Because the E3 ligase catalyzes proximity-induced ubiquitination of substrate, we questioned whether the formation of ternary complex induced by ZL216 contributed to nucleolin ubiquitination. To this end, the ubiquitination levels of nucleolin were measured. As expected, ZL216 significantly increased the ubiquitination levels of nucleolin, whereas DMSO and Control exhibited no effect on nucleolin ubiquitination (Figure 5D and 5E). Collectively, these results suggest that ZL216 promotes nucleolin ubiquitination by inducing nucleolin-ZL216-VHL ternary complex formation.

**Figure 5.**
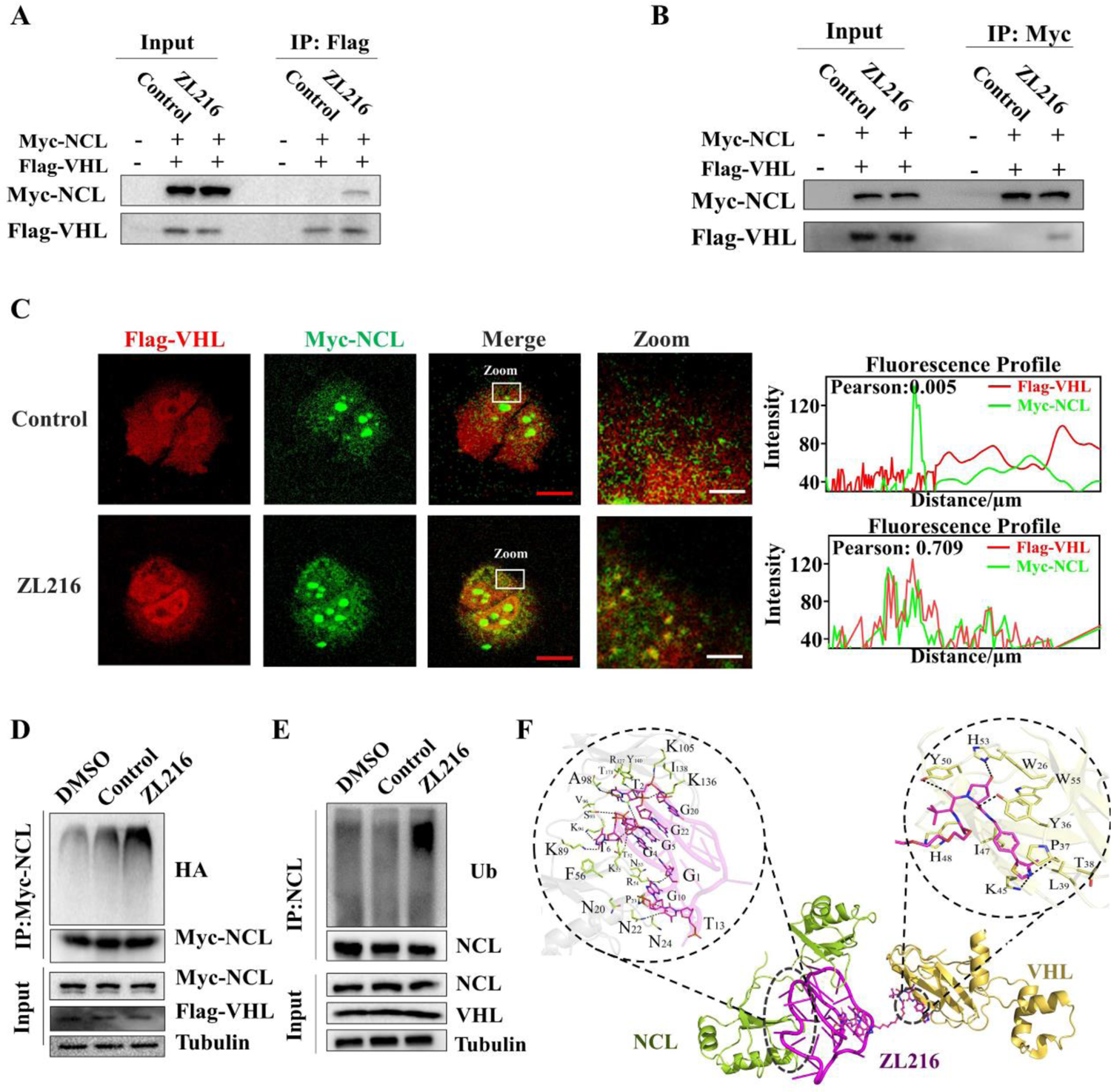
ZL216 mediates nucleolin-ZL216-VHL ternary complex formation. (A, B) MCF-7 cells transfected with plasmids expressing Flag-VHL and Myc-NCL were treated by 25 nM ZL216 or Control for 5 h. Total cell lysates were subjected to immunoprecipitation with anti-Flag (A) or anti-Myc (B) antibody. The immuno-precipitates were then analyzed by Western blotting using the indicated antibodies. (C) MCF-7 cells transfected with the indicated plasmids were treated with 25 nM ZL216 or Control for 5 h. The subcellular localization of VHL and nucleolin was visualized using immunofluorescence with the indicated antibodies. (D) MCF-7 cells transfected with plasmids expressing Myc-NCL and HA-Ub were incubated with MG132 (20 μM) for 1 h, followed by treatment with DMSO, Control (25 nM) or ZL216 (25 nM) for 5 h. Myc-NCL was immunoprecipitated with anti-Myc antibody, and the immunoprecipitates were probed with the indicated antibodies. (E) MCF7 cells were incubated with MG132 (20 μM) for 1 h, followed by treatment with DMSO, Control (25 nM) or ZL216 (25 nM) for 5 h. Nucleolin was immuno-precipitated with an anti-nucleolin antibody, and the immunoprecipitates were probed with the indicated antibodies. (F) Predicting optimal conformation of NCL: ZL216: VHL ternary complex. Light green dashes indicate the NCL structure, and blue cartoons are VHL E3 ligase. Magenta dashes represent ZL216.

To investigate the molecular basis of nucleolin-ZL216-VHL ternary complex formation, molecular docking and dynamics simulation were performed. A total of 100 conformations were extracted and ranked by docking energy value. Since low-energy conformations help reveal detailed intermolecular interaction and stable bonding interface, conformations with the lowest binding energy were chosen to predict the binding spots. The result showed that the binding energy between nucleolin and AS1411 was −123 kcal mol^-1^, whereas the docking energy between VHL and AHPC was −7.99 Kcal mol^-1^(Figure 5F). The above results implied that ZL216 could interact with nucleolin and VHL with high binding affinity.

### ZL216 inhibits the proliferation and migration of breast cancer cells via VHL binding

To address whether the ZL216-induced degradation of nucleolin affects the proliferation and migration of breast cancer cells, we performed a cell proliferation and migration assay in MCF-10A and MCF-7 cells. The results showed that ZL216 treatment dramatically inhibited the proliferation of MCF-7 cells, but had no effect on MCF-10A cells (Figure 6A and 6B). A wound healing assay demonstrated that ZL216 treatment significantly reduced the migration of MCF-7, but not MCF-10A cells (Figure 6C and 6D). Consistently, a similar result was observed in transwell assay. MCF-7, but not MCF-10A, cells in lower layer after ZL216 treatment were notably fewer in number compared to Control treatment (Figure 6E and 6F). The selective effect was consistent with the specific degradation of nucleolin induced by ZL216 in breast cancer cells. Interestingly, the effect of ZL216 on MCF-7 cells could be abolished by pretreatment with 1 μM AHPC (Figure 6), implying that the anti-proliferation and anti-migration activities of ZL216 were dependent on VHL binding. Of note, although previous studies showed that AS1411 retained its functional activity against breast cancer, we found that AS1411 treatment failed to show any effect on MCF-7 cells. This might be explained by the lower concentration of AS1411 treatment.

**Figure 6.**
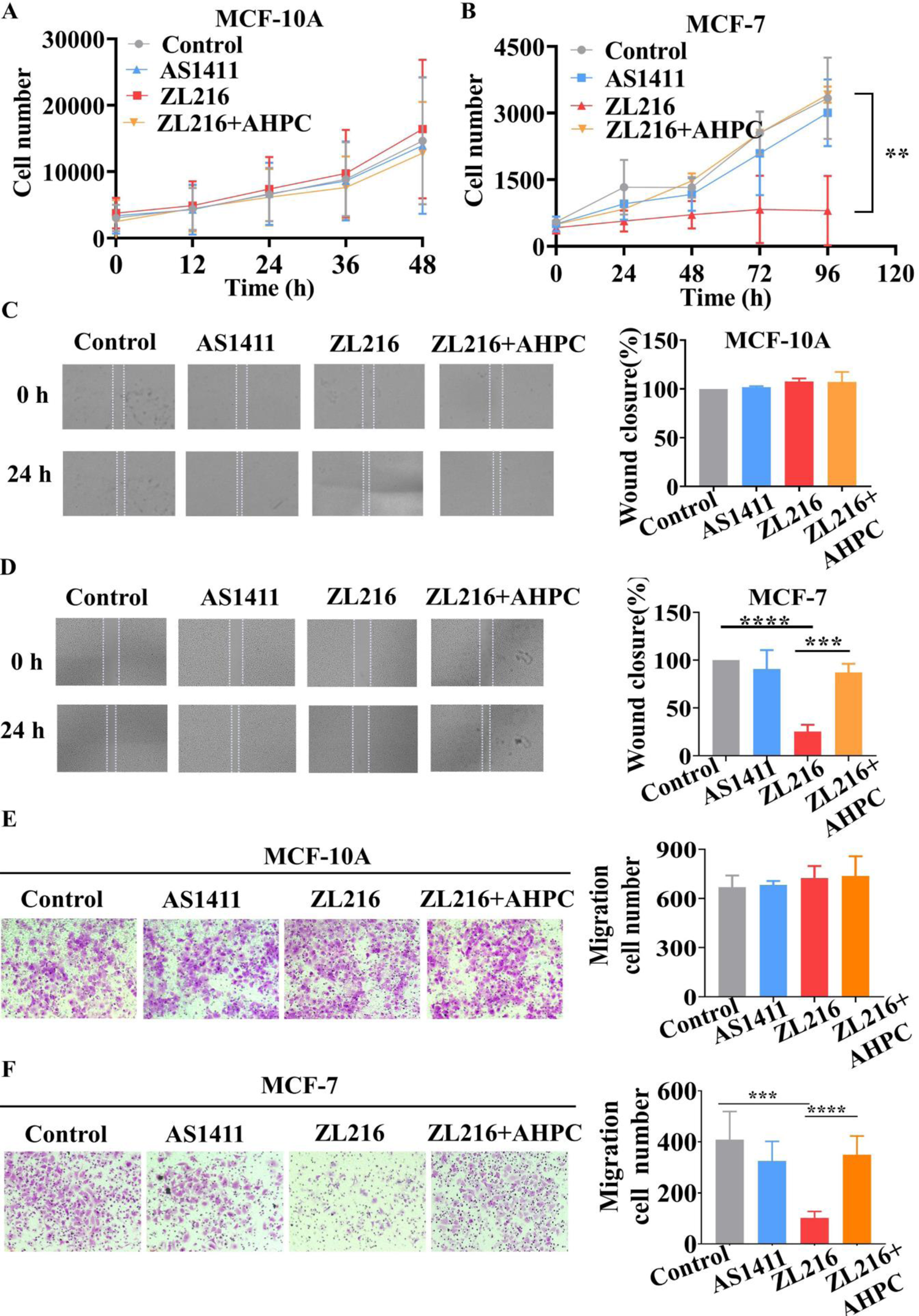
ZL216 inhibits the proliferation and migration of breast cancer cells. (A, B) cells were incubated, or not, with 1 μM AHPC for 1 h, followed by the treatment with ZL216, AS1411 or Control at the concentration of 50 nM for MCF-10A (A) or 25 nM for MCF-7 (B). Cell number was calculated at the indicated time points. (C, D) cells were incubated, or not, with 1 μM AHPC for 1 h, followed by treatment with ZL216, AS1411 or Control at the concentration of 50 nM for MCF-10A (C) or 25 nM for MCF-7 (D). Scratches were made and wound healing was observed in the indicated time points. (E, F) cells were incubated, or not, with 1 μM AHPC for 1 h, followed by treatment with ZL216, AS1411 or Control at the concentration of 50 nM for MCF-10A (E) or 25 nM for MCF-7 (F). The cells that had migrated though the pores were stained and quantified by counting five independent visual fields. (*P<0.05, **P<0.01, ***P<0.001, ****P<0.0001). Data are presented as the means ± standard deviation, n = 3.

## DISCUSSION

PROTACs function as a designed heterobifunctional molecule to induce the degradation of unwanted protein by hijacking the intracellular ubiquitin-proteasome system. Early PROTACs were developed based on bifunctional peptides that recruited E3 ligases to degrade target proteins.(25, 26) However, peptide based PROTACs suffered from low cell permeability and, hence, poor cellular activity, hindering the broad applicability of PROTACs. (27)Subsequently, owing to the identification of small molecule-based E3 ligase ligands, such as nutlin-3a for MDM2, (28) thalidomide for CRBN(29) and VH032 for VHL, (30) a series of small molecule-based PROTACs were developed, which represents a significant advancement in PROTAC technology. In this study, we further developed the first aptamer-constructed PROTAC ZL216 through combining ssDNA aptamer AS1411 with a small-molecule ligand of VHL, which demonstrate that in addition to peptides and small molecule compounds, the nucleic acid aptamers can also be used to construct PROTACs. Unlike the recently reported aptamer-conjugated BET-PROTAC using aptamer as targeted delivery tool for PROTAC in specific cell types,(22) ZL216 functioned as a novel PROTAC to promote the formation of nucleolin-ZL216-VHL ternary complex by using AS1411 as a ligand for binding to nucleolin, which potently induced the degradation of nucleolin in breast cancer cells *in vitro* and *in vivo*. Currently, few aptamers that specifically bind to E3 ligases are available; therefore, we only utilized the aptamer as a protein of interest (POI) ligand to construct PROTAC. With the development of specific aptamers for E3 ligases in the future, it will contribute to the generation of heterobifunctional aptamers to achieve the function of PROTACs.

It is estimated that the human genome encodes more than 600 E3 ligases.(30) However, the number of E3 ligases currently used for PROTAC development is still very small, mainly including VHL, CRBN, MDM2 and cIAP1.(27) Previous studies show that PROTACs based each of these E3 ligases hold different characteristics, such as specificity, efficacy, or toxicity. For example, despite using the same BCR-ABL ligand as a warhead, the CRBN-based PROTACs destroyed both BCR-ABL and c-ABL, while the VHL-based PROTACs only had an effect on c-ABL.(31) In this study, we developed a VHL-recruiting PROTAC Z216 using aptamer AS1411 as a nucleolin ligand. We demonstrated that ZL216 induced nucleolin degradation via the ubiquitin proteasome pathway in a VHL-dependent manner. Nevertheless, it is worth investigating if AS1411 can be conjugated with other E3 ligases to obtain similar effects.

PROTACs promote the ubiquitination and degradation of target protein by inducing spatial proximity between an E3 ligase and target protein via the formation of a ternary complex. The spatial distance between E3 ligase and targeted protein is critical to the efficacy of PROTACs. Of note, aptamers have a length ranging from approximately 20 to 80 nucleotides. Whether aptamer length affects the formation and stability of ternary complex and the ability of a PROTAC to degrade its target are questions that need further clarification in the future.

The water solubility of PROTACs is critical to its cell permeability and efficacy. AS1411 is a synthetic 26-base guanine-rich ssDNA oligonucleotide. Previous studies have shown that the water solubility of paclitaxel could be improved by conjugating AS1411 and paclitaxel.(32) Consistent with this, although the small molecule E3 ligase ligand APHC exhibited poor water solubility, AS1411-AHPC conjugate ZL216 showed a significant enhancement in water solubility, implying that a hydrophilic nucleic acid aptamer integrated into PROTACs can improve their water solubility.

AS1411 binds to nucleolin with high affinity and specificity. Many studies have indicated that enhanced expression of nucleolin on the cell surface is restricted to tumor cells, but not normal cells, which confers a tumor-selective binding behavior to AS1411.(17) Our results showed that the aptamer-constructed PROTAC ZL216 using AS1411 as a targeting element for nucleolin still retained the same tumor-selective property as free AS1411. Previous studies have shown that AS1411 exhibited antitumor activity. Unexpectedly, a weak activity of AS1411was observed in our studies. This phenomenon may be explained in their different modes of action. ZL216 functions as a PROTAC to promote nucleolin degradation, whereas AS1411 does not affect nucleolin levels, but rather regulates the function of nucleolin. Meanwhile, the concentration range we set, was based on the activity of ZL216, which is in nanomolar range. However, AS1411 has been reported to inhibit the growth of cancer cells in micromolar range.(30) This finding is consistent with previous studies that PROTACs generally achieve a desired pharmacological effect at significantly lower concentration owing to its catalytic mode of action.(33, 34) In addition, although we demonstrated that ZL216 could effectively degrade nucleolin *in vitro* and *in vivo* and inhibited the proliferation and migration of breast cancer cells *in vitro*, the therapeutical potential and the pharmacokinetic properties of ZL216 *in vivo* still need further evaluate.

In summary, we developed a novel PROTAC ZL216 using aptamer AS1411 directly combined with VHL E3 ligand to induce the degradation of nucleolin *in vitro* and *in vivo* by promoting the formation of a nucleolin-ZL216-VHL ternary complex in breast cancer cells, but not in normal breast cells. As a result, ZL216 inhibited the proliferation and migration of breast cancer cells. Our work proves that in addition to peptides and small molecule compounds, nucleic acid aptamers can be used to conjugate with E3 ligand to construct PROTACs and provides a promising strategy for development of tumor-selective PROTACs.

## MATERIALS AND METHODS

### Reagents

The following reagents were used in this study: the Von Hippel-Lindau (VHL) E3 ligand (S,R,S)-AHPC-PEG3-azide (AHPC) (Sigma Aldrich), MG132 (Sigma Aldrich), MLN4924 (S7109, Selleck Chemicals), anti-nucleolin antibody (D4C7O, Cell Signaling Technology), anti-β-Tubulin antibody (9F3, Cell Signaling Technology), anti-mouse IgG Alexa Fluor® 488 Conjugate (4408, Cell Signaling Technology), anti-rabbit IgG Alexa Fluor® 594 Conjugate antibody (8889, Cell Signaling Technology), anti-VHL antibody (sc-135657, Santa Cruz), anti-Ub antibody (sc-166553, Santa Cruz), anti-β-Actin antibody (AT0001, Engibody), anti-Flag antibody (20543-1-AP, Proteintech) anti-MYC antibody (M192-3, MBL International), anti-HA antibody (M180-3, Medical & Biological Laborato), Cell Counting Kit-8 (CCK-8) (New Cell & Molecular Biotech), Agarose and Super GelRedTM (S2001, US Everbright®Inc) and Cell Culture flasks (NEST Biotechnology). All DNA sequences were synthesized by Hippobio Technology (Zhejiang, China) and Shenggong Biotechnology (Shanghai, China).

### Synthesis of ZL216 and Control

(S, R, S)-AHPC-PEG3-azide was dissolved in DMSO at a concentration of 10 mM. DBCO-labeled AS1411 was dissolved in ddH2O at a concentration of 100 µM. After that, they were mixed under mechanical stirring at room temperature, and according to previous research, the click reaction between an azide and DBCO was performed overnight. The solution was then purified by HPLC and verified by mass spectrometry. The synthetic methods and identification of Control, Cy5-ZL216 and Cy5-Control were the same as described above.

### Cell Culture

Human breast cancer cell line BT474 was cultured in RPMI-1640 containing 10% fetal bovine serum (FBS). Human breast cancer cell line MCF-7 was cultured in high-glucose Dulbecco’s Modified Eagle’s Medium (DMEM) containing 10% fetal bovine serum (FBS). Normal human epithelial mammary cell line MCF-10A was maintained in complete medium purchased from Cellcook Biotech Company. All cells were cultured at 37 °C in an incubator containing 5% CO2.

### Flow Cytometry Analysis

The binding abilities of ZL216 and AS1411 were determined by flow cytometry. Cy5-labeled ZL216, AS1411 and Control were incubated with 0.2% EDTA-treated cells (2×10^5^) in 200 µL binding buffer (BB; DPBS, 5 mM MgCl2, 4.5 mg/mL glucose, 1 mg/mLBSA, and 0.1 mg/mL yeast tRNA) at 37 °C for 40 min. Then, cells were washed twice with washing buffer (WB; 5 mM MgCl2 and 4.5 g/L glucose in Dulbec-co’s PBS), resuspended in 400 µL DPBS and then subjected to flow cytometric analysis. The Control, ZL216 or AS1411 were used at the following concentration (50 nM for MCF-10A and BT474 cells, and 25 nM for MCF-7 cells) as above reported for the flow cytometry analysis.

To determine the binding site between AS1411 and ZL216, 200 nM Cy5-labeled ZL216 was severally incubated with target cells in 200 μL BB at 37℃ for 30 min, then added 200 nM,1000 nM,4000 nM of AS1411 or Control for another 30 min. Then washing cells twice with WB, the fluorescence intensity was analyzed by flow cytometry.

To explore the expression of nucleolin on the cell plasma membrane, 0.2% EDTA-treated cells were blocked in 5% BSA buffer for 30 min at 4℃ and then incubated with anti-nucleolin primary antibody (1:400 in 1% BSA buffer) for 1 h at 4 °C. Cells were washed 3× in DPBS and incubated with Alexa Fluor 647-conjugated goat anti-rabbit IgG (diluted 1:200 in 1% BSA buffer) for 30 min at room temperature. After that, cells were washed 3× in DPBS and then subjected to analysis by flow cytometry.

### Live Cell Confocal Microscopy

After 2×10^5^ cells were seeded in a 35-mm glass bottom dish for 24 h, we incubated cells with Cy5-labeled ZL216 or Control in 200 µL BB at 37℃ for 2 h. After washing cells three times with WB, confocal laser scanning microscopy (Carl Zeiss) was used to detect fluorescent signals. MCF-10A cells were treated with the 50 nM concentrations of Control, ZL216 or AS1411, while the concentration of Control, ZL216 or AS1411 in MCF-7 cells was 25 nM.

### *In Vitro* Stability Assay

2 µM Cy5-labeled AS1411 and ZL216 were incubated in human serum at 37 °C for 0, 2, 4, 6, 8, 10, 12, 24, 48, 72 and 96 h. Then, aptamers were visualized on a 0.5% agarose gel (containing 1% SDS), the band density was assayed by molecular imaging (Bio-Rad).

### Real-Time PCR

ZL216, Control, AHPC, AS1411 or DMSO treated cells at the concentration of 25 nM for MCF-7 or 50 nM for BT474 for 5 h, and the total RNA was extracted using Eastep (Promega catalog no. LS1040). 1 μg of purified RNA was reverse-transcribed in a 20 µL reaction using a FastQuant RT Super Mix Kit (TIANGEN, catalog no. KR106-02). Real-Time PCR was then performed by using Maxima SYBR Green/ROX kit (ThermoFisher Scientific) on an Applied Biosystem ABI 7500 Real-Time PCR system with the following primer sequences: NCL-F GCACCTGGAAAACGAAAGAAGG; NCL-R GAAAGCCGTAGTCGGTTCTGT; GAPDH-F ATGTTCGTCATGGGTGTGAA; GAPDH-R TGTGGTCATGAGTCCTTCCA.

### Western Blot Analysis

After 2×10^5^ cells were seeded in a 12-well plate for 24 h, we treated cells with ZL216/Control/AS1411/AHPC at certain times. Then cells were lysed in RIPA buffer (50 mM Tris (pH 7.4), 150 mM NaCl, 1% Triton X-100, 1% sodium deoxycholate, 0.1% SDS, sodium orthovanadate, sodium fluoride, EDTA, leupeptin, catalog no. P0013B, Beyotime) supplemented with proteasome inhibitors (Bimake no. B14011). Lysates were centrifuged at 15,000 rpm for 15 min, and the supernatants were used for Western blot analysis.

### Cycloheximide (CHX) Assay

MCF-7 or BT474 cells were pretreated with cycloheximide (100 μg/mL) for 1 h, prior to adding either vehicle (DMSO) or ZL216. At the indicated time points, cells were lysed. The protein levels were assessed using Western blot analysis. The concentration of ZL216 used in MCF-7 or BT474 cells was 25 nM or 50 nM respectively.

### Proteasome-Dependency Experiment

2×10^5^ MCF-7 or BT474 cells were cultured in a 12-well plate for 24 h and then incubated in 20 µM MG132 (dissolved in DMSO) or equal volume of DMSO for 1 h. Cells were then treated with Control or ZL216 for another 5 h, harvested, followed by analysis of lysates by Western blot assay. The Control and ZL216 were used at the concentration of 50 nM for MCF-10A and BT474 cells, and 25 nM for MCF7 cells.

### Co-Immunoprecipitation (Co-IP)

MCF-7 cells were transfected with Myc-NCL and Flag-VHL expression plasmids by using LipoMAXTM Reagent (SUDGEN). After 24 h, cells were treated with 20 µM MG132 for 1 h, followed by incubation with 25 nM ZL216 or Control for 5 h. Thereafter, cells were lysed in M-PER (Thermo Fisher Scientific catalog no.78501) supplemented with proteasome inhibitors (Bimake no. B14011). Cell lysates were centrifuged at 15,000 rpm for 15 min, and supernatants collected. 3 μg of relevant primary antibodies were incubated with the lysate (total proteins) at 4 °C overnight, followed by the addition of 30 µL of magnetic beads conjugated with protein G (Invitrogen, catalog no. 10004D) for 1 h and eluted by co-IP washing buffer. Finally, the pulled down proteins were dissolved in 2×SDS sample buffer, boiled, and subjected to Western blot analysis.

### *In Vivo* Ubiquitylation Assay

MCF-7 cells cotransfected, or not, with Myc-NCL, Flag-VHL and HA-ubiquitin plasmid. After 24 h, cells were pretreated with 20 μM MG132 for 1 h, followed by incubation with 25 nM ZL216 or Control for another 5 h. Cells were lysed, and lysates were then incubated with anti-Myc or an anti-NCL antibody at 4 °C overnight followed by protein A/G agarose beads for an additional 1 h at room temperature. The proteins were released from beads by boiling the beads in SDS/PAGE sample buffer. The ubiquitination of Myc-NCL or NCL was detected by Western blot with anti-Ub or anti-HA antibody.

### Immunofluorescence Staining

MCF-7 cells were cotransfected with Myc-NCL and Flag-VHL plasmids. After 24 h, cells were treated with 25 nM ZL216 for 5 h, then fixed with 4% paraformaldehyde for 15 min and washed three times with DPBS. Cells were permeabilized with 0.2% Triton X-100 for 10 min before blocking in 5% BSA for 1 h. Thereafter, cells were incubated with the primary antibodies at 4 °C overnight. Subsequently, cells were washed and labeled using secondary antibodies conjugated to Alexa Fluor 488 or Alexa Fluor 633 (Thermo Fisher) at room temperature for 1 h. Finally, a confocal microscope (Zeiss LSM510) was used to acquire fluorescent images.

### Structure determination and model building

We firstly constructed the structure of nucleolin ligand. BLAST comparison was performed on the primary structure sequence of nucleolin with all the data in the PDB database, we selected the crystal structure of nucleolin containing the corresponding DNA fragment for subsequent analysis. The structure of AS1411 was found in the RCSB PDB crystal database, and a total of 100 phases for interaction between AS1411 and nucleolin were identified by Rosetta software. Rosetta’s own clustering analysis module was used for clustering analysis and ranking according to the molecular docking energy score. The structure with most clustering and lower scores is used for subsequent analysis. Then, we performed molecular docking of VHL and AHPC ligand. VHL was flexibly butted with AHPC by using auto dock software, and the preliminary butted phase structure was obtained. The phase with the lowest docking energy was selected for structure extraction and used for subsequent molecular dynamics research. Finally, the dynamics simulation of Nucleolin-AS1411-AHPC-VHL complex system structure was performed in NAMD software with the simulation time being over 70 ns.

### Cell Migration Assay

MCF-7 and MCF-10A cells were seeded in a 12-well plate to grow until creating 95%-100% confluent monolayers. The monolayers were scratched with a sterile pipette tip and washed with serum-free medium. For MCF-7 cells, the remaining cells were treated with 25 nM ZL216, AS1411 or Control in serum-free medium. For MCF-10A cells, the remaining cells were pretreated with mitomycin C (10 μg/mL) for 2 h to block proliferation, followed by the addition of 50 nM ZL216, AS1411 or control in the complete medium. Images from the same wounded fields were obtained at 0 and 24 h under BioTek Imaging (USA). The wound healing area (area not covered by cells) was calculated using ImageJ software (NIH, Bethesda, MD, USA).

12 well-transwell chambers (Transwell, Corning Costar) were used to perform transwell migration assays. Briefly, 1×10^4^ cells were plated in the upper transwell chamber and incubated with ZL216, AS1411 or Control in serum-free medium at the concentration of 50 nM for MCF-10A cells or 25 nM for MCF-7 cells. Then, 0.5 mL complete medium was added in the lower chamber as chemoattractant. After 24 h of incubation, migrated cells on the membrane of the chambers were fixed with 4% paraformaldehyde at room temperature. After 15 min, staining with 0.1% crystal violet was performed, followed by destaining with PBS solution. The number of migrating cells was counted under a microscope in five predetermined fields.

### In Vivo Assay

Five-week-old female athymic BALB/c (BALB/c-nude) mice were used for *in vivo* fluorescence imaging and protein degradation assays. Nude mice were subcutaneously transplanted with 8×10^7^ MCF-7 cells or BT474 cells. 100 μL DPBS containing 50 μM Cy5-labeled ZL216, AS1411 or control were injected intravenously through the tail vein. The fluorescence images of live mice were collected using the IVIS Lumina II (Caliper LifeScience, USA). For the *in vivo* degradation experiment, the dissected tumor tissues were determined by Western blot assays as described above. The animal-related experimental procedures were approved by the Animal Care and Use Committee of Hunan University (SYXK 2018-0006).

## Supporting information

Supplementary Figure 1-9 will be used for the link to the file on the preprint site.

## SUPPLEMENTAL INFORMATION

The Supporting Information is available at

## ACKNOWLEDGEMENT

This work was supported by grants from the National Key Research and Development Program of China (2021YFA0909403), the National Natural Science Foundation of China (21890744 and 81672760), the Hunan Provincial Key Research and Development Plan (2018SK2128) and the Changsha Science and Technology Project (kq2001012).

## AUTHOR CONTRIBUTIONS

L. Zhang, L. Li, X. Wang contributed equally to this work.

## REFERENCES

1. Churcher, I. (2018) Protac-Induced Protein Degradation in Drug Discovery: Breaking the Rules or Just Making New Ones? J Med Chem, 61, 444–452.

2. Khan, S., He, Y., Zhang, X., Yuan, Y., Pu, S., Kong, Q., Zheng, G. and Zhou, D. (2020) PROteolysis TArgeting Chimeras (PROTACs) as emerging anticancer therapeutics. Oncogene, 39, 4909–4924.

3. Paiva, S.L. and Crews, C.M. (2019) Targeted protein degradation: elements of PROTAC design. Curr Opin Chem Biol, 50, 111–119.

4. Posternak, G., Tang, X., Maisonneuve, P., Jin, T., Lavoie, H., Daou, S., Orlicky, S., Goullet de Rugy, T., Caldwell, L., Chan, K. et al. (2020) Functional characterization of a PROTAC directed against BRAF mutant V600E. Nat Chem Biol, 16, 1170–1178.

5. Raina, K., Lu, J., Qian, Y., Altieri, M., Gordon, D., Rossi, A.M., Wang, J., Chen, X., Dong, H., Siu, K. et al. (2016) PROTAC-induced BET protein degradation as a therapy for castration-resistant prostate cancer. Proc Natl Acad Sci U S A, 113, 7124–7129.

6. Khan, S., Zhang, X., Lv, D., Zhang, Q., He, Y., Zhang, P., Liu, X., Thummuri, D., Yuan, Y., Wiegand, J.S. et al. (2019) A selective BCL-X(L) PROTAC degrader achieves safe and potent antitumor activity. Nat Med, 25, 1938–1947.

7. Hu, J., Hu, B., Wang, M., Xu, F., Miao, B., Yang, C.Y., Wang, M., Liu, Z., Hayes, D.F., Chinnaswamy, K. et al. (2019) Discovery of ERD-308 as a Highly Potent Proteolysis Targeting Chimera (PROTAC) Degrader of Estrogen Receptor (ER). J Med Chem, 62, 1420–1442.

8. Mullard, A. (2021) Targeted protein degraders crowd into the clinic. Nat Rev Drug Discov, 20, 247–250.

9. Cotton, A.D., Nguyen, D.P., Gramespacher, J.A., Seiple, I.B. and Wells, J.A. (2021) Development of Antibody-Based PROTACs for the Degradation of the Cell-Surface Immune Checkpoint Protein PD-L1. J Am Chem Soc, 143, 593–598.

10. Maneiro, M.A., Forte, N., Shchepinova, M.M., Kounde, C.S., Chudasama, V., Baker, J.R. and Tate, E.W. (2020) Antibody-PROTAC Conjugates Enable HER2-Dependent Targeted Protein Degradation of BRD4. ACS Chem Biol, 15, 1306–1312.

11. Pillow, T.H., Adhikari, P., Blake, R.A., Chen, J., Del Rosario, G., Deshmukh, G., Figueroa, I., Gascoigne, K.E., Kamath, A.V., Kaufman, S. et al. (2020) Antibody Conjugation of a Chimeric BET Degrader Enables in vivo Activity. ChemMedChem, 15, 17–25.

12. Dragovich, P.S., Pillow, T.H., Blake, R.A., Sadowsky, J.D., Adaligil, E., Adhikari, P., Chen, J., Corr, N., Dela Cruz-Chuh, J., Del Rosario, G. et al. (2021) Antibody-Mediated Delivery of Chimeric BRD4 Degraders. Part 2: Improvement of In Vitro Antiproliferation Activity and In Vivo Antitumor Efficacy. J Med Chem, 64, 2576–2607.

13. Nimjee, S.M., White, R.R., Becker, R.C. and Sullenger, B.A. (2017) Aptamers as Therapeutics. Annu Rev Pharmacol Toxicol, 57, 61–79.

14. Ye, M., Hu, J., Peng, M., Liu, J., Liu, J., Liu, H., Zhao, X. and Tan, W. (2012) Generating aptamers by cell-SELEX for applications in molecular medicine. Int J Mol Sci, 13, 3341–3353.

15. Jiang, F., Liu, B., Lu, J., Li, F., Li, D., Liang, C., Dang, L., Liu, J., He, B., Badshah, S.A. et al. (2015) Progress and Challenges in Developing Aptamer-Functionalized Targeted Drug Delivery Systems. Int J Mol Sci, 16, 23784–23822.

16. Wu, X., Liu, H., Han, D., Peng, B., Zhang, H., Zhang, L., Li, J., Liu, J., Cui, C., Fang, S. et al. (2019) Elucidation and Structural Modeling of CD71 as a Molecular Target for Cell-Specific Aptamer Binding. J Am Chem Soc, 141, 10760–10769.

17. Reyes-Reyes, E.M., Teng, Y. and Bates, P.J. (2010) A new paradigm for aptamer therapeutic AS1411 action: uptake by macropinocytosis and its stimulation by a nucleolin-dependent mechanism. Cancer Res, 70, 8617–8629.

18. Soundararajan, S., Chen, W., Spicer, E.K., Courtenay-Luck, N. and Fernandes, D.J. (2008) The nucleolin targeting aptamer AS1411 destabilizes Bcl-2 messenger RNA in human breast cancer cells. Cancer Res, 68, 2358–2365.

19. Cheng, Y., Zhao, G., Zhang, S., Nigim, F., Zhou, G., Yu, Z., Song, Y., Chen, Y. and Li, Y. (2016) AS1411-Induced Growth Inhibition of Glioma Cells by Up-Regulation of p53 and Down-Regulation of Bcl-2 and Akt1 via Nucleolin. PLoS One, 11, e0167094.

20. Rosenberg, J.E., Bambury, R.M., Van Allen, E.M., Drabkin, H.A., Lara, P.N., Jr., Harzstark, A.L., Wagle, N., Figlin, R.A., Smith, G.W., Garraway, L.A. et al. (2014) A phase II trial of AS1411 (a novel nucleolin-targeted DNA aptamer) in metastatic renal cell carcinoma. Invest New Drugs, 32, 178–187.

21. Soundararajan, S., Wang, L., Sridharan, V., Chen, W., Courtenay-Luck, N., Jones, D., Spicer, E.K. and Fernandes, D.J. (2009) Plasma membrane nucleolin is a receptor for the anticancer aptamer AS1411 in MV4-11 leukemia cells. Mol Pharmacol, 76, 984–991.

22. He, S., Gao, F., Ma, J., Ma, H., Dong, G. and Sheng, C. (2021) Aptamer-PROTAC Conjugates (APCs) for Tumor-Specific Targeting in Breast Cancer. Angew Chem Int Ed Engl, 60, 23299–23305.

23. Pichiorri, F., Palmieri, D., De Luca, L., Consiglio, J., You, J., Rocci, A., Talabere, T., Piovan, C., Lagana, A., Cascione, L. et al. (2013) In vivo NCL targeting affects breast cancer aggressiveness through miRNA regulation. J Exp Med, 210, 951–968.

24. Fonseca, N.A., Rodrigues, A.S., Rodrigues-Santos, P., Alves, V., Gregório, A.C., Valério-Fernandes, Â., Gomes-da-Silva, L.C., Rosa, M.S., Moura, V., Ramalho-Santos, J. et al. (2015) Nucleolin overexpression in breast cancer cell sub-populations with different stem-like phenotype enables targeted intracellular delivery of synergistic drug combination. Biomaterials, 69, 76–88.

25. Sakamoto, K.M., Kim, K.B., Kumagai, A., Mercurio, F., Crews, C.M. and Deshaies, R.J. (2001) Protacs: chimeric molecules that target proteins to the Skp1-Cullin-F box complex for ubiquitination and degradation. Proc Natl Acad Sci U S A, 98, 8554–8559.

26. Schneekloth, J.S., Jr., Fonseca, F.N., Koldobskiy, M., Mandal, A., Deshaies, R., Sakamoto, K. and Crews, C.M. (2004) Chemical genetic control of protein levels: selective in vivo targeted degradation. J Am Chem Soc, 126, 3748–3754.

27. Pettersson, M. and Crews, C.M. (2019) PROteolysis TArgeting Chimeras (PROTACs) - Past, present and future. Drug Discov Today Technol, 31, 15–27.

28. Schneekloth, A.R., Pucheault, M., Tae, H.S. and Crews, C.M. (2008) Targeted intracellular protein degradation induced by a small molecule: En route to chemical proteomics. Bioorg Med Chem Lett, 18, 5904–5908.

29. Fischer, E.S., Böhm, K., Lydeard, J.R., Yang, H., Stadler, M.B., Cavadini, S., Nagel, J., Serluca, F., Acker, V., Lingaraju, G.M. et al. (2014) Structure of the DDB1-CRBN E3 ubiquitin ligase in complex with thalidomide. Nature, 512, 49–53.

30. Buel, G.R., Chen, X., Chari, R., O’Neill, M.J., Ebelle, D.L., Jenkins, C., Sridharan, V., Tarasov, S.G., Tarasova, N.I., Andresson, T. et al. (2020) Structure of E3 ligase E6AP with a proteasome-binding site provided by substrate receptor hRpn10. Nat Commun, 11, 1291.

31. Lai, A.C., Toure, M., Hellerschmied, D., Salami, J., Jaime-Figueroa, S., Ko, E., Hines, J. and Crews, C.M. (2016) Modular PROTAC Design for the Degradation of Oncogenic BCR-ABL. Angew Chem Int Ed Engl, 55, 807–810.

32. Li, F., Lu, J., Liu, J., Liang, C., Wang, M., Wang, L., Li, D., Yao, H., Zhang, Q., Wen, J. et al. (2017) A water-soluble nucleolin aptamer-paclitaxel conjugate for tumor-specific targeting in ovarian cancer. Nat Commun, 8, 1390.

33. Zou, Y., Ma, D. and Wang, Y. (2019) The PROTAC technology in drug development. Cell Biochem Funct, 37, 21–30.

34. Schapira, M., Calabrese, M.F., Bullock, A.N. and Crews, C.M. (2019) Targeted protein degradation: expanding the toolbox. Nat Rev Drug Discov, 18, 949–963.

