## Supplementary Figure 1-9 will be used for the link to the file on the preprint site. for "Development of a novel PROTAC using the nucleic acid aptamer as a targeting ligand for tumor selective degradation of nucleolin"

**Supplemental Information**

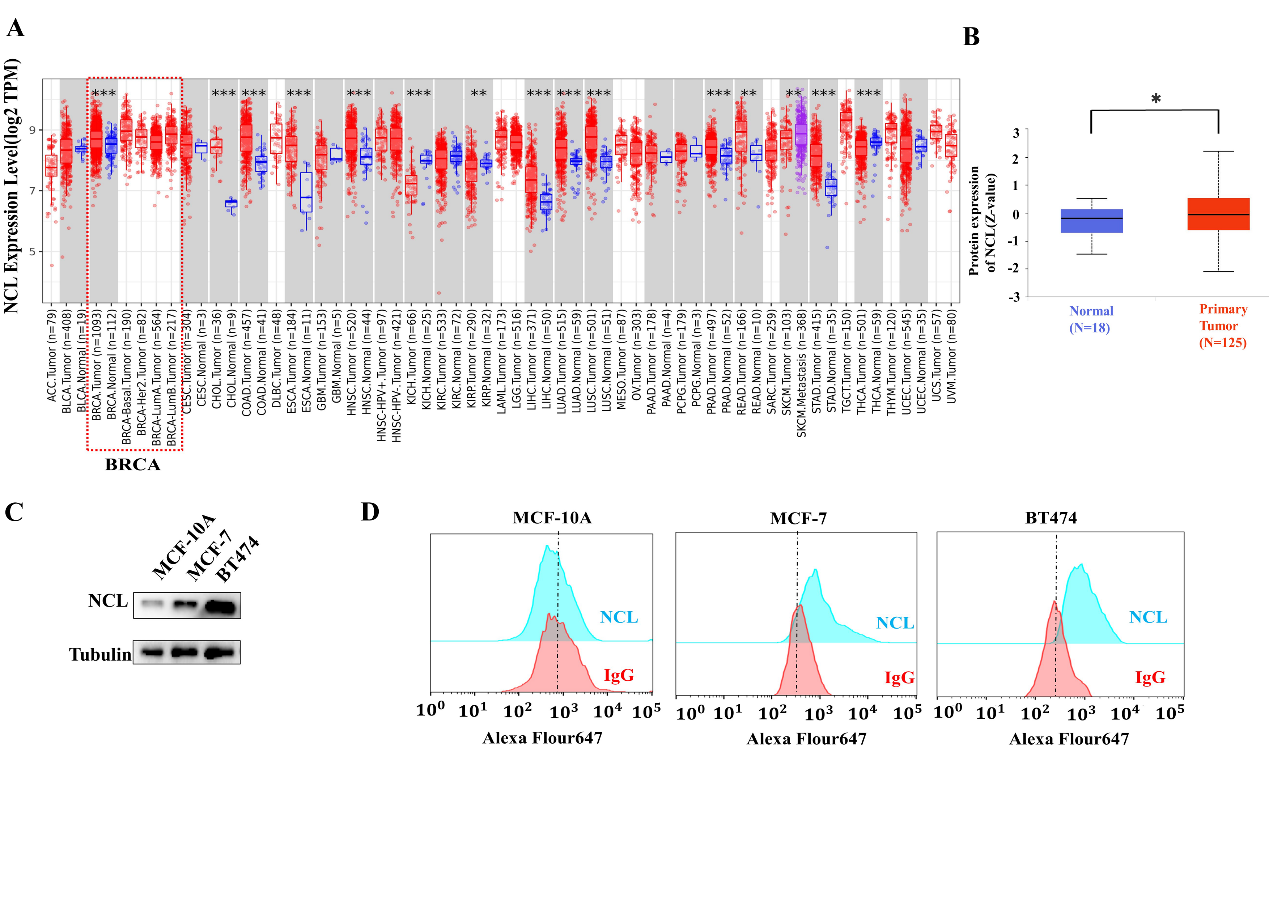

**Figure S1. Expression of nucleolin (NCL) in tumors.** (A) Human nucleolin expression levels in different tumor types from the TCGA database were determined by TIMER2.0. (*P<0.05, **P<0.01, ***P<0.001). (B) The expressions of nucleolin protein in normal breast and breast cancer tissues were analyzed using CPTAC dataset (*P< 0.05). (C) Total protein from MCF-10A, MCF-7 and BT474 cells was extracted and subjected to Western blotting using the indicated antibodies. (D) MCF-10A, MCF-7 and BT474 cells were incubated with anti-nucleolin antibody or control IgG. The expression of nucleolin on cell membranes was analyzed by flow cytometry.

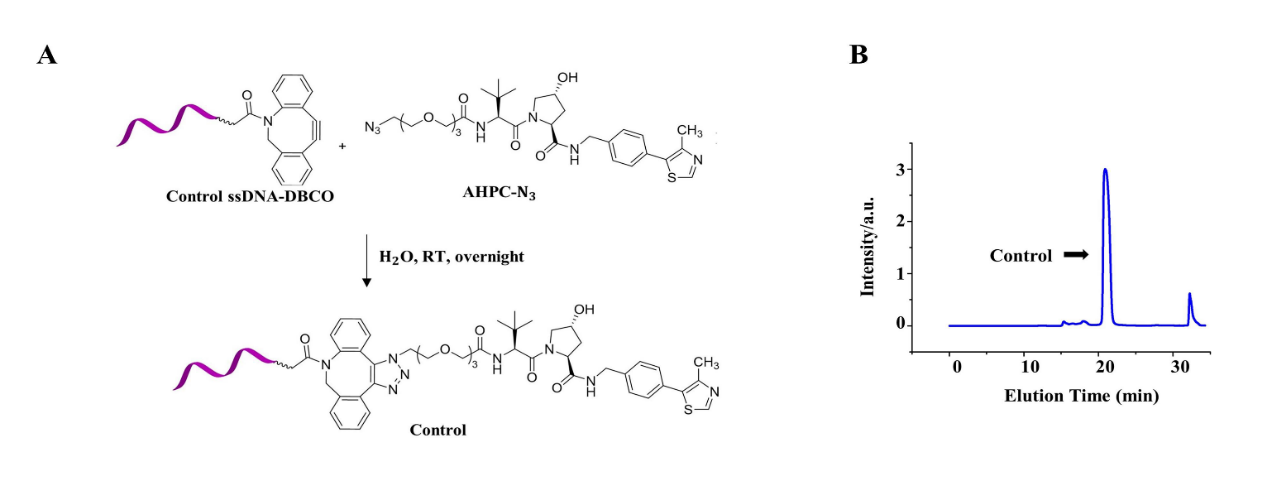

**Figure S2. The synthesis of control**. (A) Synthetic roadmap of Control. (B) The purification of Control by HPLC.

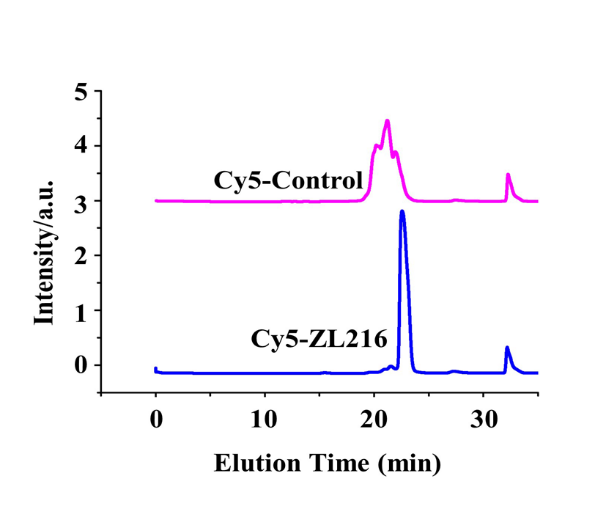

**Figure S3.** The purification of Cy5-labeled Control and ZL216 by HPLC.

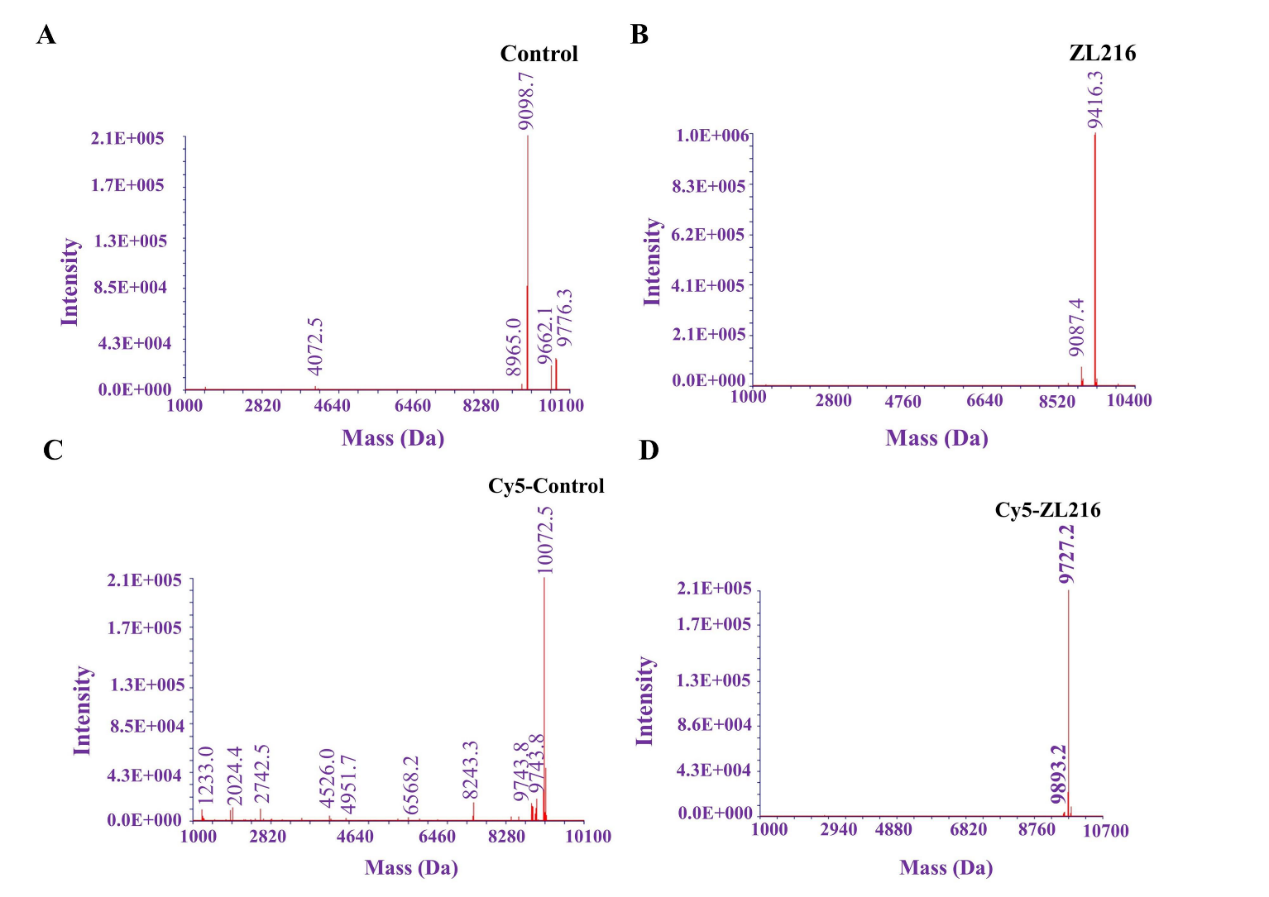

**Figure S4. The LC-MS analysis of conjugates.** Control (A), ZL216 (B), Cy5-Control (C) and Cy5-ZL216 (D) were purified by HPLC and analyzed by DNA mass spectrometry.

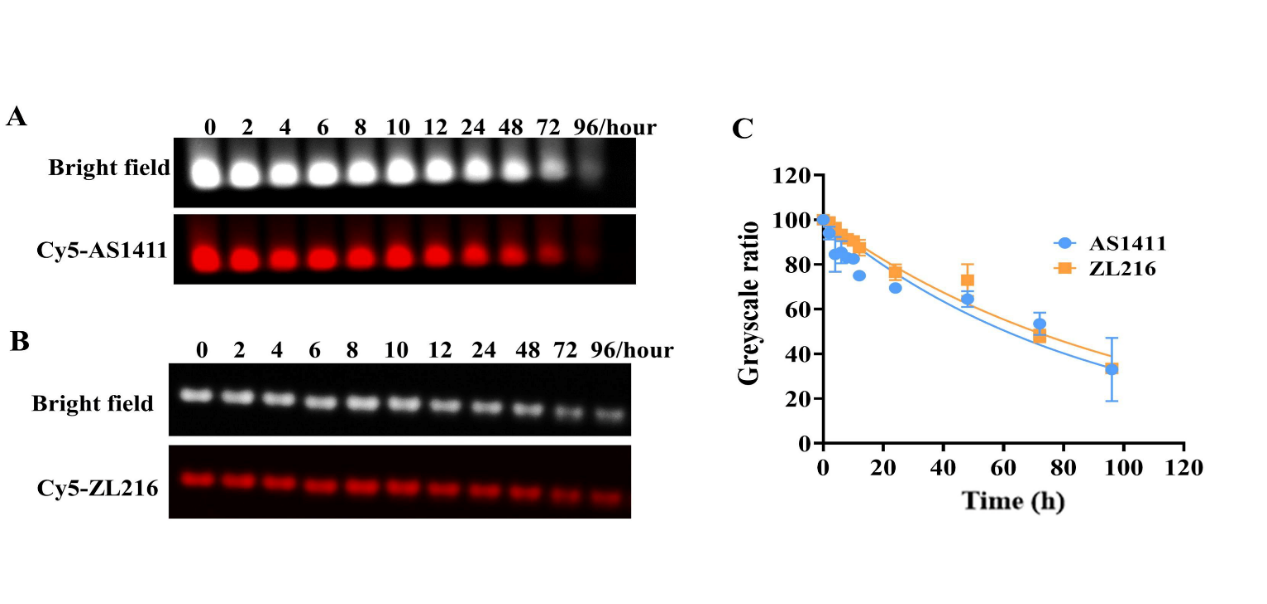

**Figure S5. The stability of AS1411 and ZL216 in serum.** (A-B) Agarose gel electrophoresis was employed to analyze the contents of AS1411(A) and ZL216 (B) after incubation with serum for the indicated times. (C) The levels of AS1411 and ZL216 were quantified after incubation with serum for the indicated times. Data are presented as the means ±SD, n = 3.

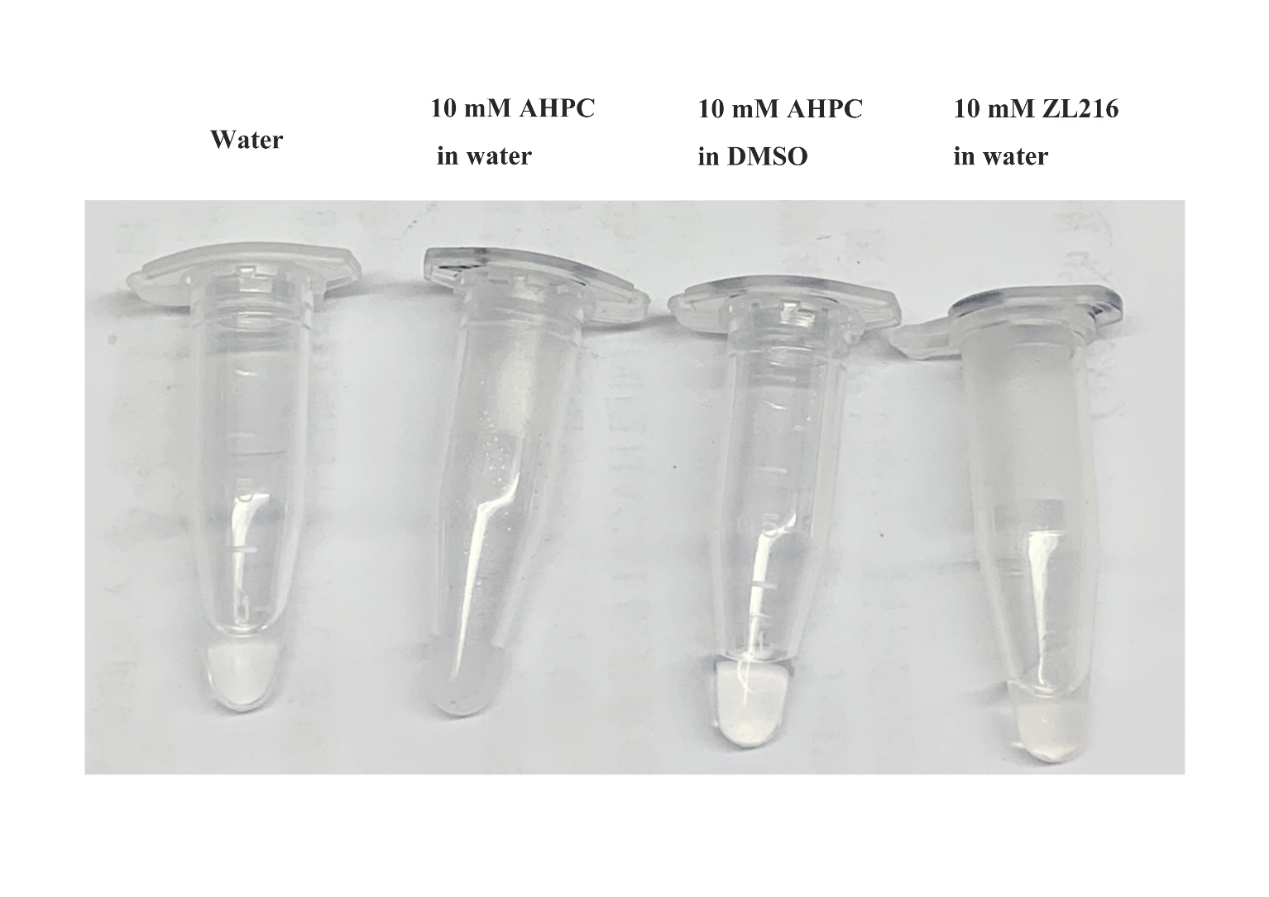

**Figure S6.** Water solubility of free AHPC and ZL216. Solubility of AHPC in water is no more than 10 mM, while the solubility of ZL216 in water is at least greater than 10 mM.

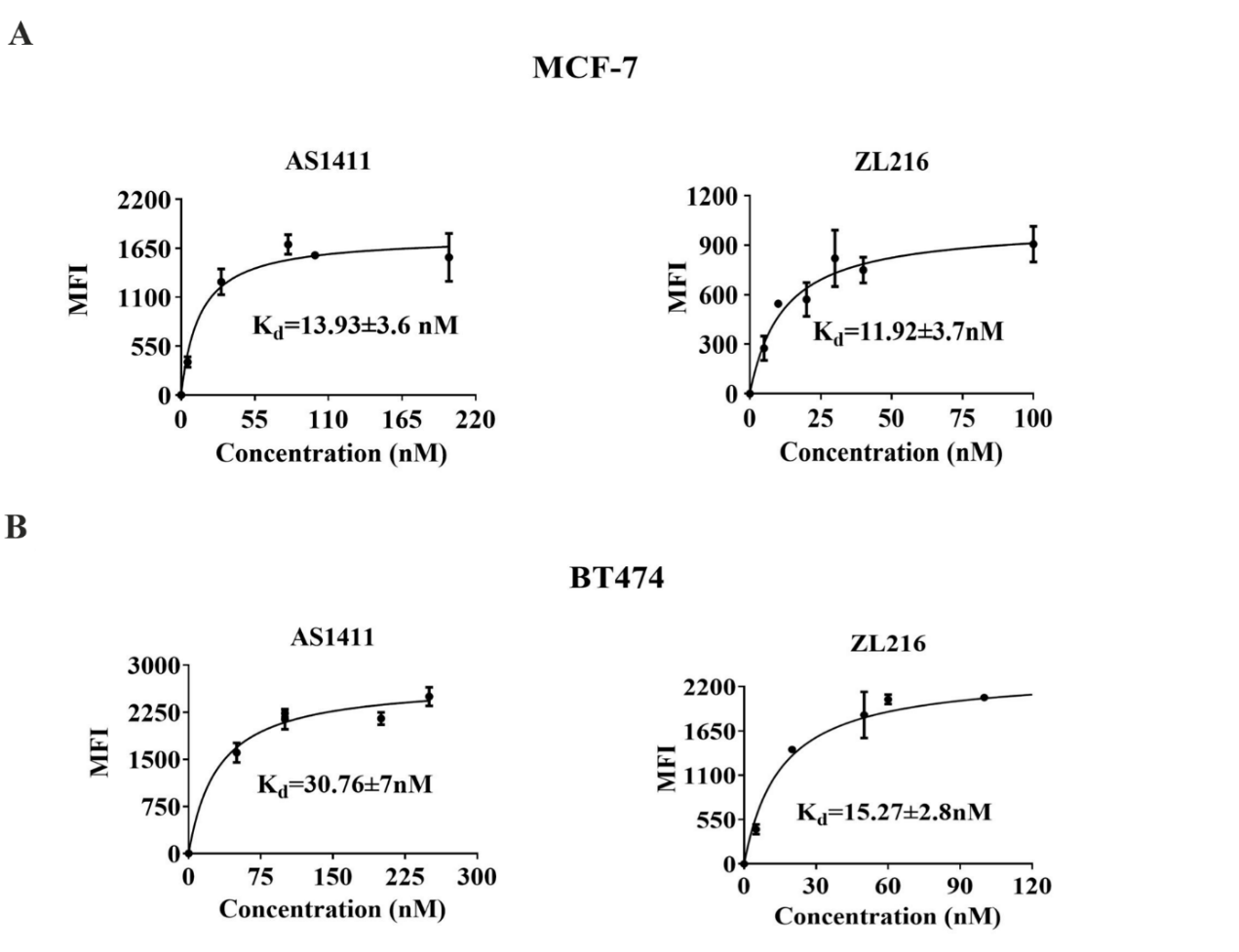

**Figure S7. The binding affinity of AS1411 and ZL216 to breast cancer cells.** (A-B) The dissociation constant of AS1411 and ZL216 for MCF-7 (A) and BT474 (B) cells was determined by flow cytometry (Median fluorescence intensity, MFI). Data are presented as the means ±SD, n = 3.

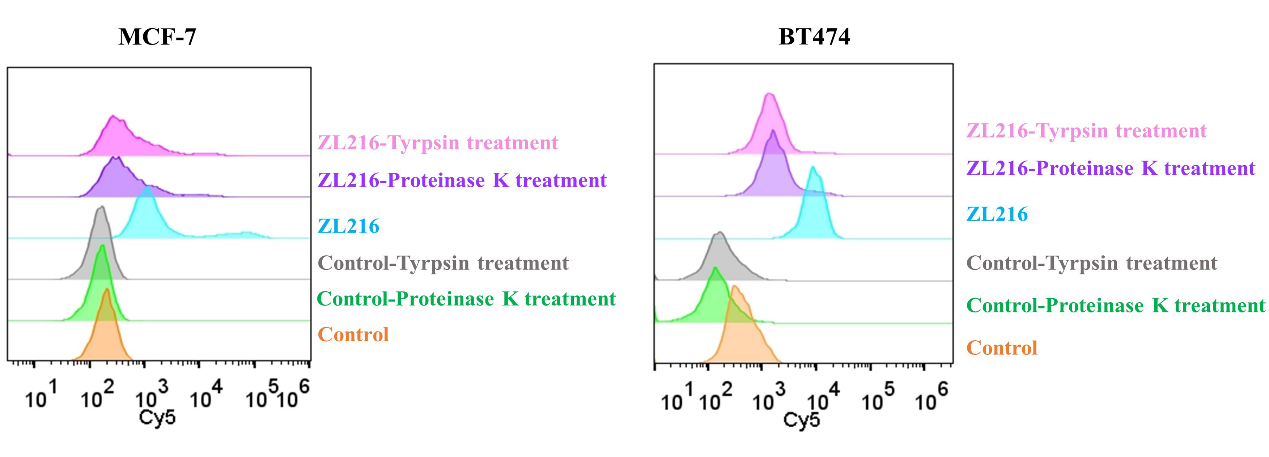

**Figure S8.** **The internalization of ZL216 into breast cancer cells.** (A-B) Cy5-modified ZL216 or Control treated MCF-7 cells (A) at the concentration of 25 nM or BT474 cells (B) at the concentration of 50 nM for 1 h at 37°C, followed by treatment with trypsin or proteinase K. The binding was analyzed by flow cytometry.

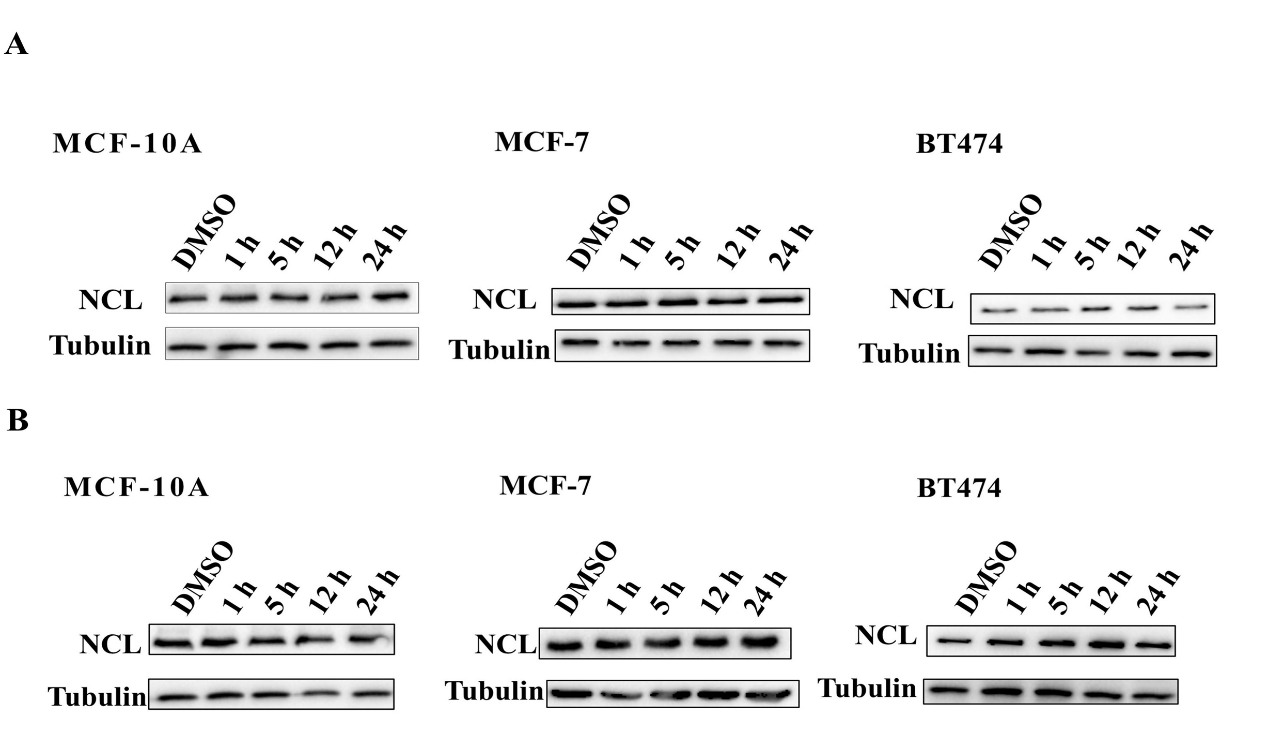

**Figure S9.** **The effect of AS1411 and AHPC on nucleolin levels.** AS1411 (A) or AHPC (B) treated cells at the concentration of 50 nM for MCF-10A, 25 nM for MCF-7 and 50 nM for BT474. Total protein was extracted at the indicated time points and subjected to Western blotting using the indicated antibodies.
